# Multinational qEEG developmental surfaces

**DOI:** 10.1101/2019.12.20.883991

**Authors:** Shiang Hu, Ally Ngulugulu, Jorge Bosch-Bayard, Maria L. Bringas-Vega, Pedro A. Valdes-Sosa

## Abstract

The quantitative electroencephalogram (qEEG) is a diagnostic method based on the spectral features of the resting state EEG. The departure of spectral features from normality is gauged by the *z* transform with respect to the age-adjusted mean and deviation of normative databases – known as the developmental equations/surfaces. However, the extent to which the data collected from different countries with various equipment require separate developmental equations remains unanswered. Here, we analyzed the EEG of 535 subjects from 3 countries, Switzerland, the USA and Cuba. The EEG power spectra of all samples were log transformed and their relations to the covariables (‘age’, ‘frequency’, ‘country’ and ‘individual’) were analyzed using the linear mixed effects model. We found that the origin ‘country’ of the subjects did not play a significant effect on the log spectra, even without interactions with other independent variables, whereas, ‘age’ and ‘frequency’ were highly significant. To estimate the developmental surfaces in greater detail, we carried out kernel regression (lowess) in two dimensions of log-age and frequency. We found two main phenomena: 1) slow rhythms (*δ, θ*) predominated in the lower ages and then decreased with a tendency to disappear at higher ages; 2) *α* rhythm was absent at lower ages, but gradually appeared more relevant in occipital and parietal regions, and increased with aging with an increasing centering frequency of *α* rhythm. We consider both phenomena as an expression of healthy neurodevelopmental and maturation related to age. It is the first study of multinational qEEG developmental surfaces accounting for ‘country’. The results demonstrate the possibility of creating international qEEG norms since the ‘individual’ and ‘age’ variability are much larger than the specific factors like ‘country’, or the technology employed ‘device’.

## Introduction

As a powerful tool, the electroencephalogram (EEG) has been widely applied to noninvasively study the brain functions in neurological and psychiatric populations (Cohen 2017). The power spectra summarize stationary and linear characteristics of EEG activities and is useful to characterize different behavioral states. Resting state EEG spectral patterns consist of the four fundamental rhythmical bands (Babiloni 2018): the slow frequency (0.5-3.5Hz) high amplitude δ rhythm associated with sleep, the fasting in frequency (3.5-7.5Hz) and increasing in amplitude θ rhythm related with drowsiness condition, the *α* (8-12Hz) rhythm associated with relaxing and eyes-closed condition, and *β* (13-30Hz) rhythm in an awake, alert state. Specially, the *α* rhythms peaks reflect the performance in various cognitive functions (John et al. 1988). The power spectra have therefore been widely used to evaluate information processing and brain states (Babiloni et al. 2004).

The quantitative EEG (qEEG) is a diagnostic method based on the extraction of spectral features from the resting state EEG. The age-dependent mean and standard deviation of a normative database is defined as the ‘developmental equations’ by (John et al. 1980) in order to complement the visual inspection of EEG and facilitate an objective evaluation of the main EEG features as the descriptive parameters from raw EEG recordings. The descriptive parameters are rescaled by the *z* transform but with the problem that the parameters are highly age dependent. Moreover, the descriptive parameters use broadband spectra that estimate on a frequency band rather individual frequency bin. Univariate *z* transform of parameters defined from the normative database is as follows

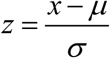

where *x* is any parameter, *μ* represents the population mean, and *σ* denotes the population standard deviation of the parameter. Age and other covariates are introduced to increase sensitivity and specificity. This was demonstrated by (John et al. 1980) and later confirmed by (Matoušek and Petersén 1973; John et al. 1977; Alvarez et al. 1987; Amador et al. 1989). Thus, normative data is usually summarized by means of age regression functions for broadband spectra.

Alvarez compared the children from Cuba and the USA (Alvarez et al. 1987) using the developmental equations originally obtained at the USA by (John et al. 1980) who argued that the norms are relatively independent of sociocultural variables and found transcultural validity of the norms. Note that his study was using broadband spectral parameters (BBSP). Similarly, the study compared using microstates at milliseconds level with the multinational EEG data (n=496, Switzerland, US, Cuba) confirmed the strong relation of EEG with age (Koenig et al. 2002). Unfortunately, it is irrelevant to the discussion of the frequency evolution since they didn’t conduct any form of frequency analysis. This present study is, therefore, the first to report multinational norms. One of the main findings from the Cuban group in the construction of EEG norm was the distinction between broad bands (*δ*, *θ*, *α*, *β*) and narrow bands in bins of frequency, for example, the most employed were from 0.39Hz to 19 Hz because of the technological constraints of the equipment. The history of the evolution of the qEEG modeling and the construction of EEG norms by the Cuban group is extensively described in (Hernandez-Gonzalez et al. 2011) where two stages of the Cuban human brain mapping project are explained with the first phase in the 1990s and the second between 2004-2006. High-resolution spectral models were explored as an alternative to BBSP. This was demonstrated by (Szava et al. 1994) using the EEG developmental surfaces avoiding the frequency and spatial smearing that may occur using BBSP. Receiver operator characteristic curve (ROC) analysis demonstrated the increased diagnostic accuracy of high-resolution spectral methods. As well, the regression equations were introduced by (Amador et al. 1990) to calculate developmental surfaces by characterizing the age frequency distribution of mean and standard deviation of the log power spectra in a normative sample. The normative surfaces allowed the calculation of *z* transformed spectra for all electrodes and *z* maps for each frequency bin.

In neuroscience, a difficult task is to obtain normative data from large samples to infer the neural substrate underlying different cognitive emotional and mental states in a healthy population. The norms can provide the information to identify pathological states, neurological and psychiatric disorders as well as the effectiveness of different treatments or interventions. But why is it so difficult to obtain multinational norms using neuroimaging data? Let’s take the example of research using magnetic resonance imaging (MRI). The MRI scanners are quite different and thus very complicated to unify the recordings and to ultimately make norms. Some solutions could be introduced in the equation as different gradients related to the type of scanner, recording parameters and normalization methods, but the comparison is still unreliable. This is a well-recognized problem. For the EEG equipment, we also meet a similar situation, that is to cope with the differences between amplifiers, recording systems, reference electrode problem (Hu et al. 2018a) and so on. The large international human brain projects are planning to gather thousands of subjects in longitudinal studies, but most of them have not included the EEG, with MRI playing the main role. Our intention is to use the EEG recording already stored in the Cuban human brain mapping (CHBMP) to validate our hypothesis and thus pave the way for the re-introduction of EEG as a main player in global brain projects.

The main questions are: Are the EEG narrow band characteristics different enough to create necessary independent norms by countries? Or can we construct international norms with big samples by adding more statistical power to the study of EEG?

The hypothesis is: There are no differences between countries in the EEG basic features in healthy subjects. We based the hypothesis on the first findings reported in (John et al. 1977) where is said for BBSP that the EEG could be cultural and race free instrument. This has not yet been confirmed for EEG narrowband spectra analysis.

The novelty of this study consisted of calculating the multinational norms with a large sample of healthy subjects since nobody compares before developmental surfaces of the log spectra norms across countries.

## Materials and methods

### Data Samples

We analyzed the resting state EEG recordings with 10-20 electrode placement system of 535 subjects: 162 subjects in 1990s and 88 subjects in 2004 from the two phases of the CHBMP, Cuban neuroscience center, Havana, Cuba, 43 subjects from the University Hospital of Clinical Psychiatry, Bern, Switzerland, and 242 subjects from the Brain Research Laboratory, Medical School, New York University. All the recordings were collected under equivalent conditions. The inclusion/exclusion criteria for the normal subjects have been described respectively in (John et al. 1977; Alvarez et al. 1987; Koenig et al. 2002; Hernandez-Gonzalez et al. 2011). The range of age of these recordings is 5.35-97 yrs. The distribution of ages of all the samples and the samples in the different datasets are shown in Figure 1A and Figure 1B, respectively. In Figure 1A, it shows that this multinational dataset contains more samples in the childhood and adolescence, moderate samples in adulthood with middle ages, and fewer samples > 65 yrs. In the Figure 1B, the dataset from Cuba 1990s covered 5.35-97 ages nearly the whole lifespan, the dataset from Cuba 2004 mainly covered the adulthoods from 17.5-47.22 yrs., the datasets from Switzerland and the United States are 10.17-16.25 yrs. and mainly 6.02-25.93 yrs., respectively. No gender information was available. To our concern, this was not an obstacle because our study didn’t include any hypothesis regarding sex.

**Figure 1.**
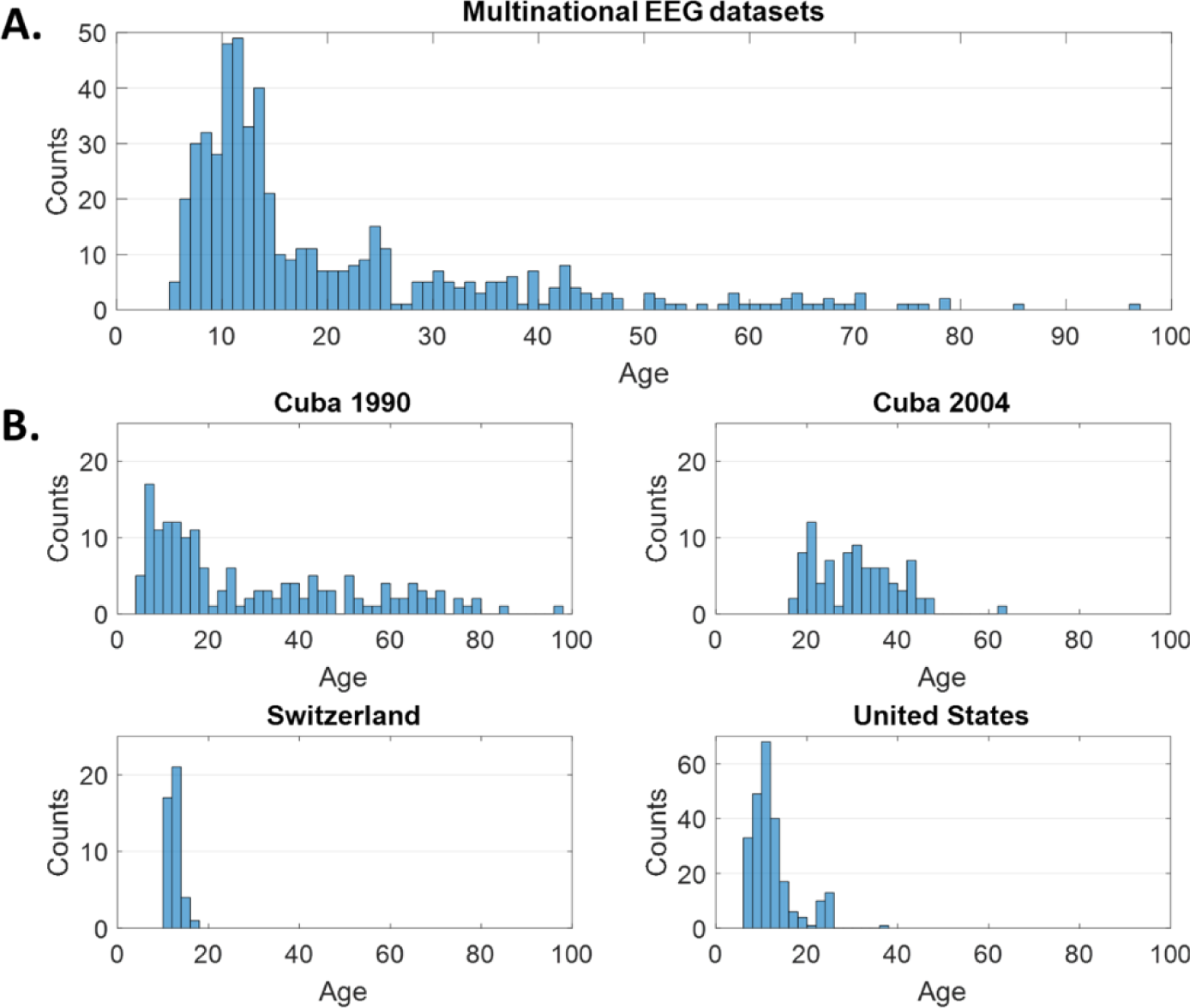
Histogram of age distribution over the lifespan in the multinational datasets. A: the histogram of age values of the total EEG datasets. B: the separate histograms of age values for different national datasets.

The Cuban subjects were a random sample of 603 subjects drawn from the total population (around 110,000) of Havana city, Cuba (Hernandez-Gonzalez et al. 2011). Eyes-closed resting digital EEG recordings were acquired with the simultaneously unipolar recording (Hu et al. 2018b, 2019) of the international 10-20 electrode placement system (Fp1, Fp2, F3, F4, C3, C4, P3, P4, O1, O2, F7, F8, T3, T4, T5, T6, Fz, Cz and Pz), with the reference to the linked earlobes. The schematic representation of this electrode layout is displayed in Figure 2 and the reference montage is shown in Figure 3B. In all the cases, the resting state EEG was recorded in a quiet, dimly light, and air-conditioned room. During the recording, the subject rested on a comfortable, half reclined armchair. Neuromeric system MEDICID 03 (NEURONIC S. A.) was employed and it is shown in Figure 3A. Note that the two phases of EEG acquisition in Cuba were using the same type of amplifier but Cuba 2004 dataset used more channels than Cuba 1990s. For the Switzerland dataset, a Nihon-Kohen EEG standard equipment was utilized (not shown here) but with similar characteristics to the Cuba equipment. To acquire the US EEG dataset, a custom-designed digital EEG data acquisition and analysis platform (DEDAAS) shown in Figure 3B was constructed by (Thatcher, R. W., & John 1977) at the Brain Research Laboratory, New York University.

**Figure 2.**
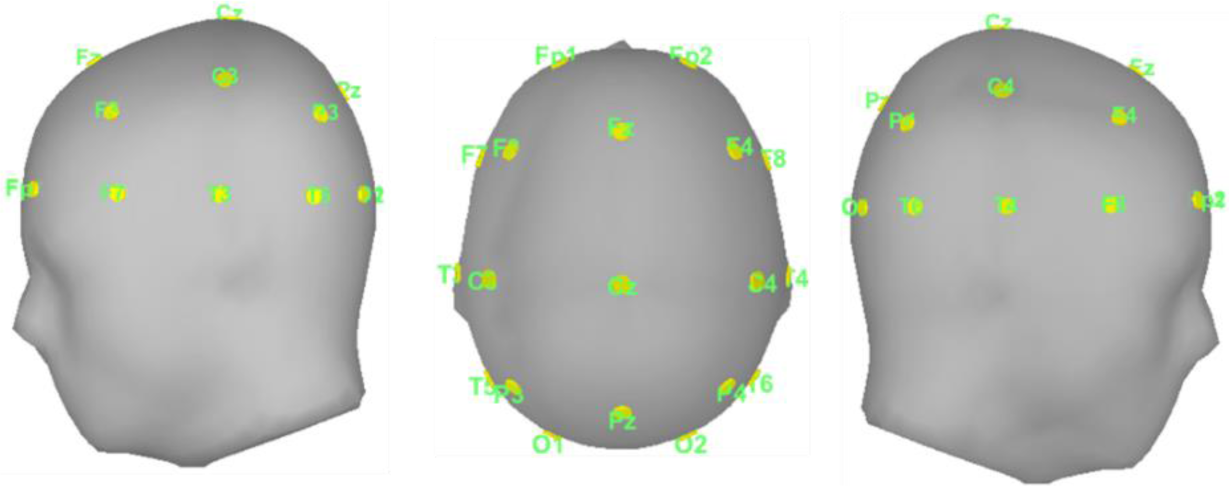
Schematic representation of the international 10-20 electrode placement system with the template from ICBM152 (Tadel et al. 2011).

**Figure 3.**
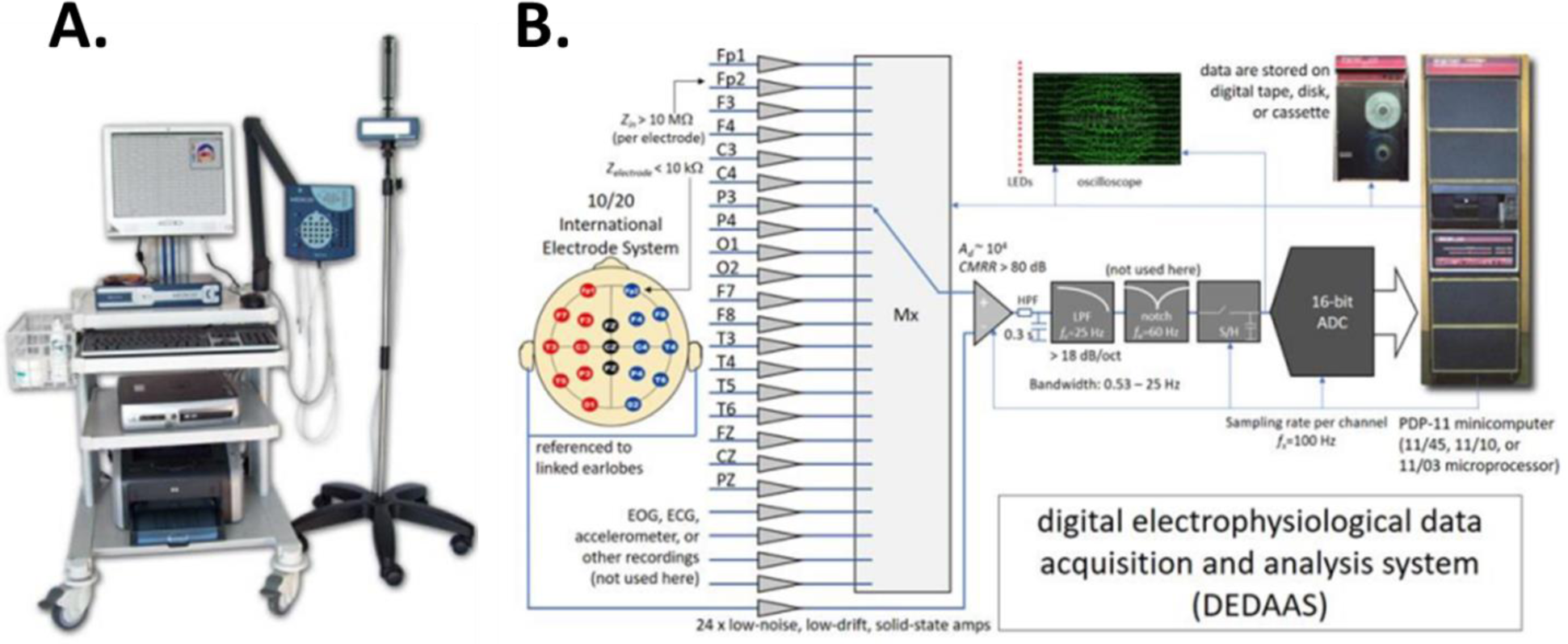
Multinational EEG acquisition systems. A: MEDICID 03 Neurometric system used in recording the datasets of Cuba 1990s and 2004 while Cuba 2004 used more channels than Cuba 1990s; B: DEDAAS in the recording the US dataset designed by E. R. John in 1970s. The Switzerland dataset was obtained from a Nihon-Kohen EEG standard equipment which was not shown here but different from the shown two.

All the recordings were acquired with the same sampling rate of 200 Hz. The minimal length of available data for each subject was at least 60 seconds of continuous resting state eyes-closed artifact-free EEG.

Informed consent was written from all participants and approvals were granted by the ethics committee of the different centers involved in the studies.

### Data analysis

We adopted the qEEG technique to identify the descriptive parameters with the steps: 1) re-referencing the selected EEG segment to the average reference; 2) estimate the scalp EEG cross-spectrum with Bartlett’s method (J. Møller 1986) by averaging the periodograms of more than 20 consecutive and non-overlapping epochs of 512 samples, regardless of possible discontinuity in the recovered records. This yielded the cross-spectra for 49 frequency bins from 0.3906Hz to 19.14Hz with a frequency resolution of 0.3906Hz. Global scale differences in scale among EEG datasets were corrected using the geometric power (Hernández et al. 1994); 3) only the diagonal entries of the cross-spectra were retained, yielding a final set of 931 (19 channels*49 frequencies) scalp power spectral features. In the present study, the cross-spectra between electrode pair is not considered; 4) the spectra at each frequency of each subject was taken the log10 transformation.

To derive the developmental surfaces with a group of variables such as ‘age’, ‘frequency’, ‘country’ and ‘individual’, we adopted the linear mixed-effects model (Demidenko 2004; McCulloch and Neuhaus 2005) that can describe the relationship between a response variable and independent variables, with coefficients varying with respect to one or more grouping variables. This model consists of fixed effects, usually the conventional linear regression part and random effects associated with individual experimental units drawn randomly from a specific regional population. The random effects have prior distributions the fixed effects do not have. A standard form of a linear mixed-effects model in this study can be expressed as

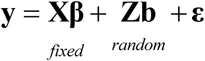

where 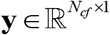 is a vector of spectra over all the electrodes*frequencies, 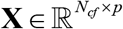 is a fixed effect design matrix, **β** ∈ ℝ^*p×1*^ is a vector of fixed effects coefficients, 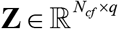 is a random effects design matrix, **b** ∈ #x211D;^*q×1*^ is a vector of random effects coefficients, and 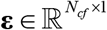 is a vector of observation residuals.

In the analysis of all the EEG datasets, the use of linear mixed-effects model enables testing the effects of ‘country’ and ‘individual’ to explain the variability of the log spectra. The MATLAB built-in function *fitlme* can fit a linear mixed effects model modeling a response variable as a linear function of fixed-effect predictors and random effect predictors. The fixed effects were ‘age’ within the range of 5.35-97 yrs. and ‘frequency’ from 0.3906Hz to 19.14Hz; and random effects are the ‘country’ set as 1, 2, 3, 4 to match ‘Cuba 2004’, ‘Cuba 1990’, ‘Switzerland’ and ‘United States’ and the ‘individual’ labeled from 1 to 535.

To select the best linear mixed effect model, we tested different models to compare between them and decided which one has the best adjustment. Using three cases, we alternatively take out one or two random effects. The ‘Wilkinson notation’ is used to explain the model fit. The three cases are:

**Case I:** Fixed effects (‘age’ and ‘frequency’), random effects (‘country’ and ‘individual’)

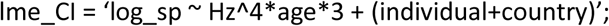

**Case II:** Fixed effects (‘age’ and ‘frequency’), random effect (‘individual’)

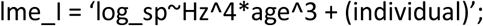

**Case III:** Fixed effects (‘age’ and ‘frequency’)

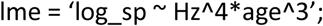

However, there is a difficulty that the singularity of the design matrix may affect the inference. To ensure the adjustment of the model after the comparison of three cases, a more complex model – the stepwise linear regression model could be used. It can help us to find the appropriate subset of regressors (independent variables) involving two opposing objectives: 1) the regression model needs to be as complete and realistic as possible. Every regressor even less related to the response variable needs to be included; 2) we need to include as few variables as possible because each irrelevant regressor decreases the precision of estimated coefficients and predicted values. The presence of extra variables increases the complexity of data collection and model maintenance. The goal of variable selection becomes one of the parsimonies: balancing between simplicity (as few regressors as possible) and fit (as many regressors as possible).

Another step was the application of kernel regression between the log10 spectra and the frequencies to see how surfaces evolve with log10 age. We employed a locally weighted scatterplot smoothing (LOWESS) method (Cleveland 1979; Cleveland and Devlin 1988) which is a nonparametric regression method that combines multiple regression models in a k-nearest-neighbor (KNN) based metamodel. It was built on classical methods, such as least square regression and address situations in which the classical procedure does not work well or cannot be effectively applied without undue labor, combining much of the simplicity of linear least square regression with the flexibility of nonlinear regression. It does this by fitting simple models to localized subsets of the data to build up a function that describes the deterministic part of the variation in the data, point by point. In fact, one of the chief attractions of this method is that the data analyst is not required to specify a global function of any form to fit a model to the data, only to fit segments of the data.

## Results

### Spectral analysis and scale removal

Below we show the EEG spectra of three subjects randomly selected but with the same age around 20 yrs. from the Cuba 1990, Switzerland and the United States as examples, Figure 4A shows different amplitudes in the log10 scale of the power spectra for the dataset from different nations. The strikingly distinct amplitudes indicate that the spectra are affected by the scale. It was thus imperative to remove the scale factor before investigating the ‘country’ effect in creating the qEEG developmental surfaces. Figure 4B is the log10 spectra after removing the scale factor by subtracting the mean log10 spectra over all the electrodes and frequencies from the original log10 spectra.

**Figure 4.**
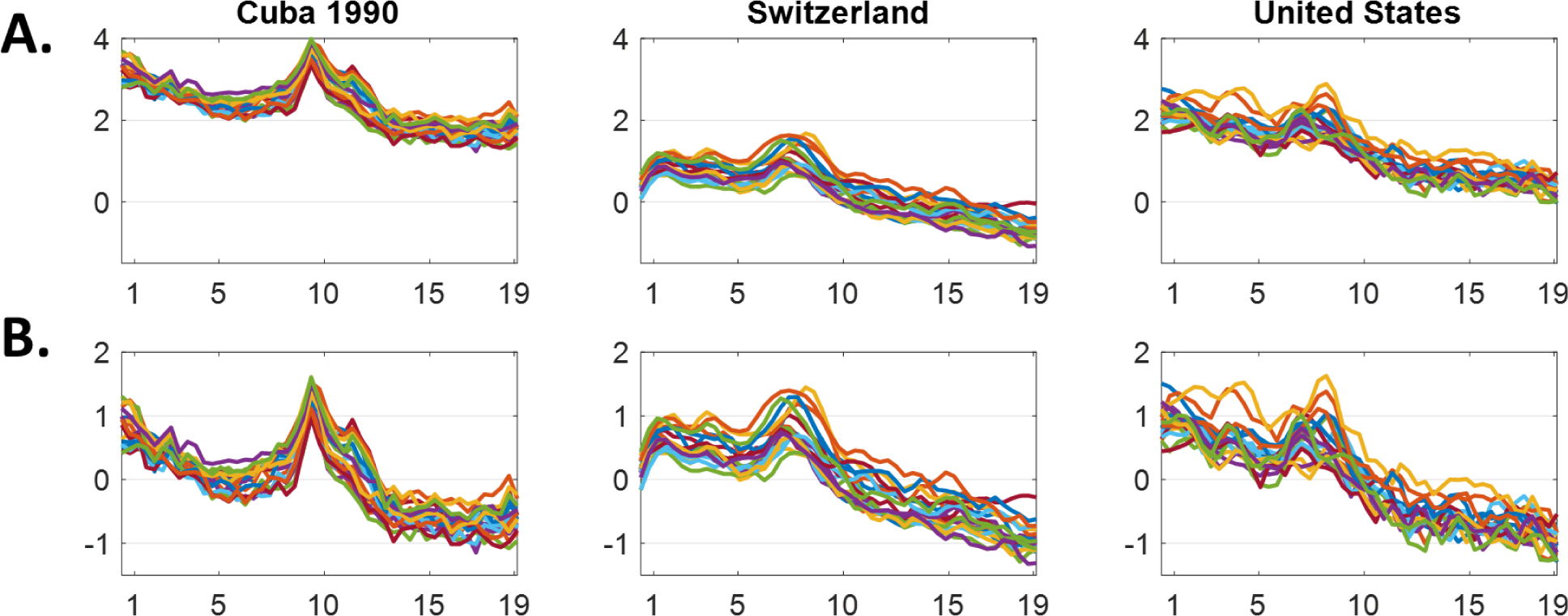
Spectra analysis and scale factor removal. A: Raw scale; B: Scaling factor removed. For all the charts, Y axes – log10 of the spectra, X axes – frequency bins from 0.3906Hz to 19.14Hz, different colors of curves – distinguishing the 19 electrodes. Cuba 2004 is not shown because it is on the same scale as Cuba 1990s.

### Linear mixed-effects model

To compare the models in the three cases described in the section ‘Data analysis’, we employed another routine *compare* (MATLAB built-in function) to compare the Case I with II, and the Case II with III using formulas: ‘result = compare(lme_CI, lme_I, ‘NSim’, ‘1000’)’ and ‘result = compare (lme_I, lme, ‘NSim’, 1000)’, where 1000 repetitions of simulations were done. Figure 5 illustrated the comparison between the models for the 19 electrodes. The comparison between the Case I and II was not significant (p = 0.08) and the comparison between the Case II and III was also no significant (p = 0.49). We concluded that the effects of ‘individual’ and ‘country’ were not important so that these two variables can be eliminated which means that the Case III (‘log_sp ~ Hz^4*age^3’) is the simplest and can fit the data.

**Figure 5.**
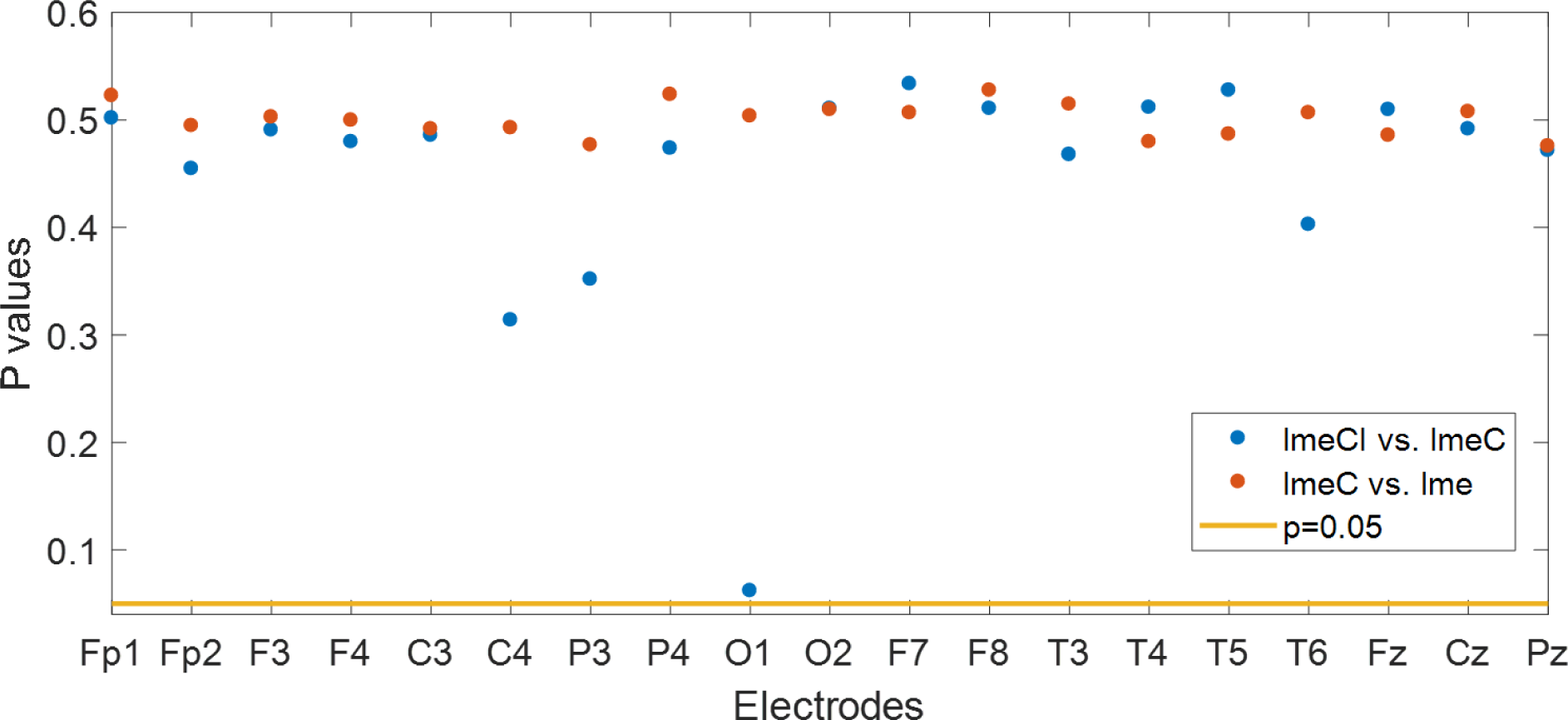
Model comparisons using the linear mixed effects model for 19 electrodes.

### The stepwise linear regression model

We selected higher order variable (polynomial power 5) because the first order is linear with only power 1 which had a fitting, none of the variables is significant using the formula to fit the model ‘log_sp ~ Hz^5*age^5’. Table 1 shows the *t-*test and *p-*value associated with the comparison of age and frequency for polynomial power 5 in the electrode O1, though the analysis with the same procedure was done for all the electrodes. You can see that the coefficients increase with the higher order of the polynomial. This indicated the complexity of the data might be explained by the higher order parametric model. For that reason, we introduce the non-parametric method – kernel regression next.

**Table 1.**
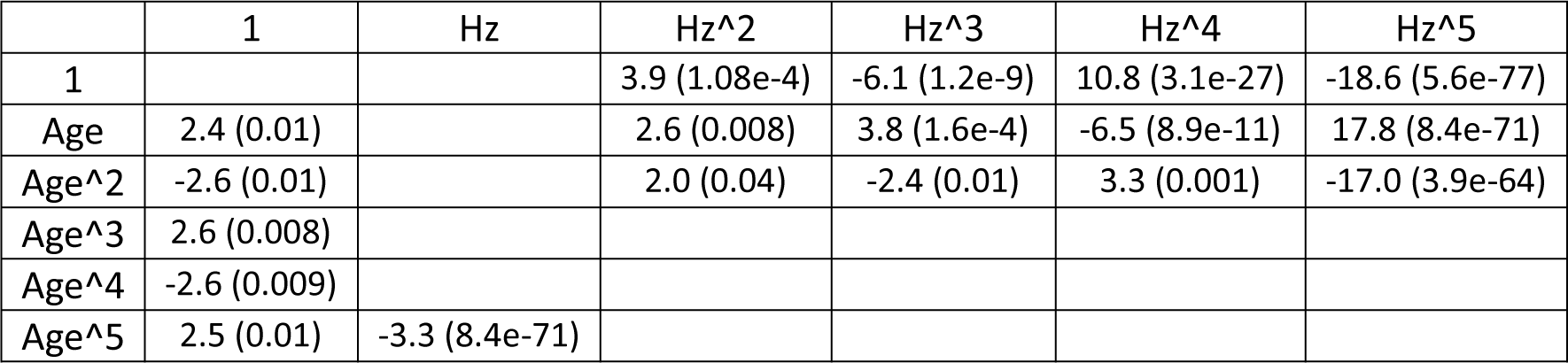
t (p) values of stepwise linear regression with the model ‘log_sp ~ Hz^5*age^5’ with the electrode O1 as an example.

|  | 1 | Hz | Hz<sup>2</sup> | Hz<sup>3</sup> | Hz<sup>4</sup> | Hz<sup>5</sup> |
| --- | --- | --- | --- | --- | --- | --- |
| 1 |  |  | 3.9 (1.08e-4) | -6.1 (1.2e-9) | 10.8 (3.1e-27) | -18.6 (5.6e-77) |
| Age | 2.4 (0.01) |  | 2.6 (0.008) | 3.8 (1.6e-4) | -6.5 (8.9e-11) | 17.8 (8.4e-71) |
| Age<sup>2</sup> | -2.6 (0.01) |  | 2.0 (0.04) | -2.4 (0.01) | 3.3 (0.001) | -17.0 (3.9e-64) |
| Age<sup>3</sup> | 2.6 (0.008) |  |  |  |  |  |
| Age<sup>4</sup> | -2.6 (0.009) |  |  |  |  |  |
| Age<sup>5</sup> | 2.5 (0.01) | -3.3 (8.4e-71) |  |  |  |  |

### Kernel regression

An important correction was to calculate the logarithm of the age to plot the results. This is because, on the one hand, the slow *δ* and *θ* rhythms have a fast decrease in the lower ages with a strong tendency to disappear with neurodevelopment and maturation; on the other hand, the *α* rhythm is growing with age, showing a different trajectory as you can see in Figure 6.

**Figure 6.**
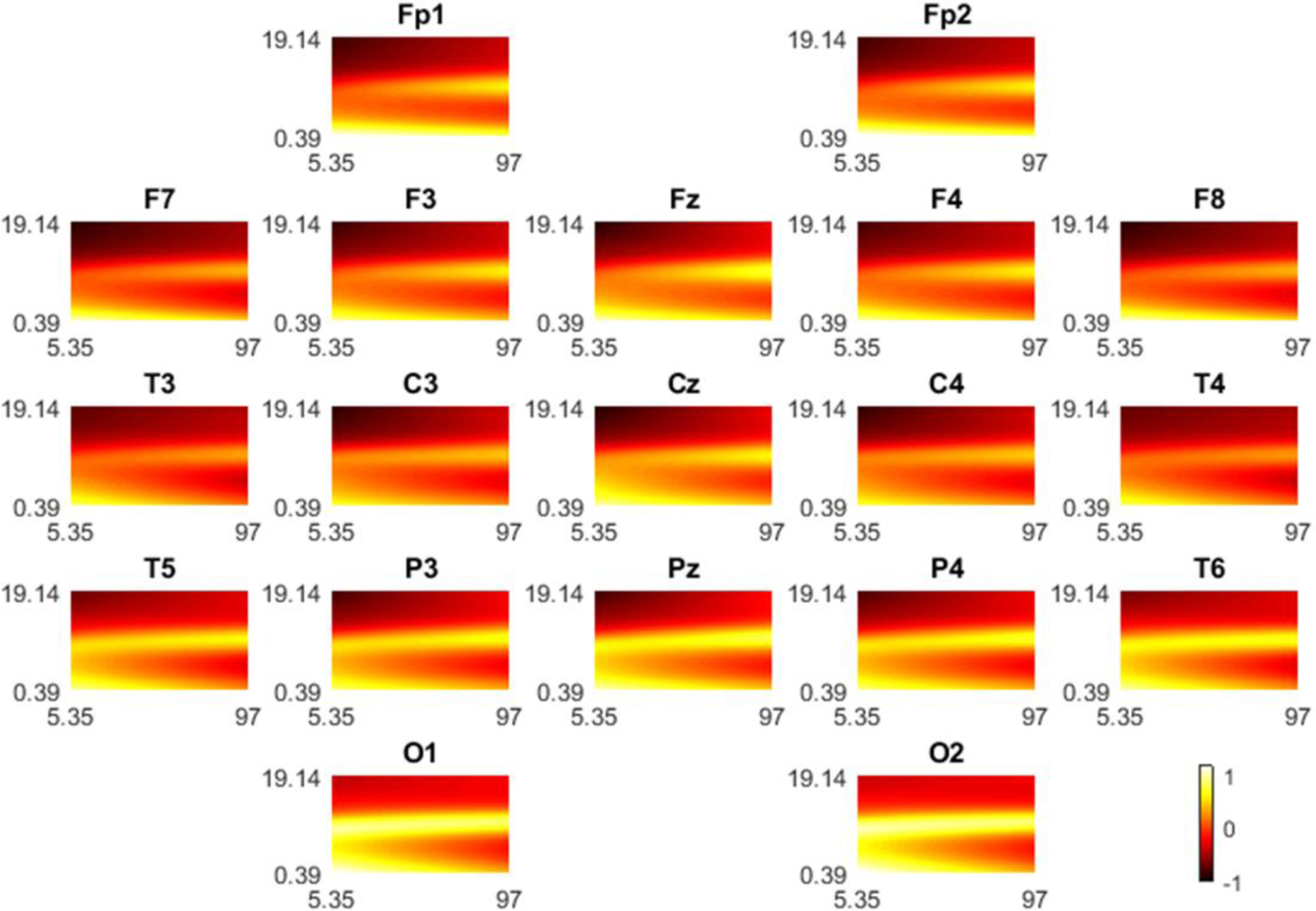
Developmental surfaces of spectra with age (X-axes: 5.35 – 97 yrs.) and frequencies (Y axes: 0.39 – 19.14 Hz). X-axes are plotted in the log10 scale; Y axes are plotted in the natural scale.

## Discussion

This study confirmed that the age of the individual subject and the EEG frequencies have highly significant relations as well as interactions with the log10 spectra of the EEG in all channels of the 10-20 system as shown in Table 1.

These results are in a good agreement with the findings from the previous research (Benninger et al. 1984; Smit et al. 2012; Vandenbosch et al. 2019) that ‘age’ plays a crucial role in the individual neurodevelopment. On the one hand, our study established that individual countries of origin do not have a significant impact on EEG development surfaces. This supports the earlier study by Alvarez compared children of Cuba and US (Alvarez et al. 1987) using the equations originally obtained at the US by (John et al. 1980) who argued that the norms were relatively independent of sociocultural influences. Therefore, it is arguable that the country of origin does not reflect the developmental surfaces (equations) describing the brain maturation.

Previously, resting state EEG power spectra had been shown to be a robust biomarker of neurodevelopment. Age-related maturation of the EEG power has long been noted, with attenuated slower (*δ* and *θ*) rhythms and increased faster (*α* and *β*) rhythms with increasing age (Ahn et al. 1980; John et al. 1980; Gasser et al. 1988; Lüchinger et al. 2012). This has been corroborated by (Lubar 1985) who reported a pattern of lower *θ* activity in the resting EEG of healthy children, with significantly higher *α* and scattered results for the *β* range (>14 Hz). In this present study, a novel and more detailed description of the developmental surfaces was obtained using a nonparametric regression technique – LOWESS. This highlighted how age affects the expression of EEG activities. As expected, the slow (*δ* and *θ*) rhythms decrease with age but *α* rhythm increased with age as indicated in Figure 6 from which it is evident how the resting state EEG spectra evolve with age.

The EEG of typically developing children shows a developmental increase in *α* rhythm and a decrease in *θ* rhythm. Concurrent resting-state fMRI and EEG measures demonstrated that EEG changes during development may reflect age-related changes in emphasis and integration of local and long-range networks, as inferred from spatial coherency in BOLD signals during resting periods (Lüchinger et al. 2012). The EEG maturational findings abovementioned have been comprehensively quantified over the lifespan with the data stored in the Cuban human brain mapping project with ages 5.35-97 yrs., with the estimation of ‘developmental equations’ for scalp EEG spectra (Szava et al. 1994) and developmental surfaces created in this present study. These developmental surfaces (equations) are the yardsticks against which we can compare the EEG of pathological populations.

Finally, the model adopted in the present study proved to be effective for creating the multinational qEEG norms using mixed-effects models. In the future, we need to test this norm in other countries with even more diversity in socio-cultural backgrounds to replicate this result. Also, we need to test the sensitivity and specify this method and move this research to calculate multinational norms by using features from EEG source analysis. An even larger challenge is to use connectivity measures derived from the source analysis.

## Conclusion

For the first time, we created international norms including countries (Cuba, US, and Switzerland) with quite different sociocultural background, and demonstrated that the EEG activity is free of the ‘country’, ‘amplifier’, and ‘individual’ effect. It is thus unnecessary to calculate specific norms for countries if the appropriate correction methods for the individuals are applied, such as removing the general scale factors in the magnitude of the spectra. A machine learning approach can be applied in the future to predict age but without the inclusion of ‘country’ and ‘individual’.

## Acknowledgment

This study was funded by the National Natural Science Foundation of China (No. 61871105, 81601585, 81861128001), the CNS program of UESTC (No. Y03111023901014005) and the 111 project B12027. The authors declare no conflicts of interest.

